# Delivery of BACE1 siRNA mediated by TARBP-BTP fusion protein reduces β-amyloid deposits in a transgenic mouse model of Alzheimer’s Disease

**DOI:** 10.1101/097121

**Authors:** Mohamed M Haroon, Kamal Saba, Venkata H Boddeda, Jerald M Kumar, Anant Bahadur Patel, Vijaya Gopal

## Abstract

Systemic delivery of nucleic acids to the central nervous system (CNS) is a major challenge for the development of RNA interference-based therapeutics due to absence of stability, target specificity, non-permeability to the blood-brain barrier (BBB), and mainly due to lack of suitable carriers. Using a designed bi-functional fusion protein TARBP-BTP, very recently we demonstrated knockdown of target genes in the brain of both AβPP-PS1 (Alzheimer’s disease, AD) and wildtype C57BL/6 mice upon systemic delivery of a single dose of siRNA. In this report, we further substantiate this hypothesis through an extended study in AβPP-PS1 mice, which upon treatment with seven doses of β-secretase APP cleaving Enzyme 1 (BACE1) siRNA, led to target-specific effects in the mouse brain. Concomitant gene silencing and consequent reduction in plaque load in the cerebral cortex and hippocampus (>60%) in mice treated with TARBP-BTP:siRNA complex further led to improvement in spatial learning and memory, which was assessed and verified through Morris Water Maze test that revealed significant improvement in cognitive function. Moreover, the treatment did not induce any adverse effects as revealed by the histopathology of different organs. The work validates the efficiency of TARBP-BTP fusion protein as an efficient mediator of RNAi giving considerable scope for future intervention of neurodegenerative disorders of the CNS through the use of short nucleic acids as gene specific inhibitors.

## Introduction

Alzheimer’s disease (AD), an age-related neurodegenerative disorder is typically characterized by death of neurons, accretion of neurofibrillary tangles containing hyperphosphorylated tau and extracellular deposition of amyloid (Aβ) plaques in the brain^1,2^. The primary molecular events following onset and pathogenesis of AD has demonstrated the major role of Aβ in disease progression particularly in brain regions involved in cognition and memory. Early pathological corroboration that Aβ, a derivative of APP, has a critical role in the disease progression^3,4^, led to the development of Aβ-related treatment protocols to inhibit the successive cleavage of β and γ-secretase, and thereby reduce the liberation of Aβ. Studies subsequent to these findings targeted the competing α-secretase in addition to the β and γ-secretase to modulate Aβ production that was viewed as a viable approach towards anti-amyloid therapy^2^.

Although there are several approaches that deliver therapeutic molecules to the brain^5^, considering the variable nature of pathogenesis of the disease at the time of diagnosis, the efficacy of these molecules in treatment are unpredictable. In addition, treatment of AD is also challenging due to the presence of the BBB comprising of endothelial cells lining the inner lumen that divides the circulating blood from the central nervous system (CNS)^6,7^, precluding effectual outcomes. Strategies involving reduction of Aβ by targeting β-secretase have shown that there are no adverse consequences of knocking down expression of this gene^8,9^. However, in many of the aforesaid strategies, the decreased efficacy is in part attributed to lack of target reach, target specificity and toxicity hindering further advancement in disease intervention.

Efforts towards delivery of drugs to the brain in the recent past have demonstrated the use of peptides and fusion proteins to deliver RNAi therapeutics to the CNS. However, the delivery of siRNA *in vivo* entails the usage of carrier molecules that are safe, biocompatible, physiologically stable and easy to administer. Through examples mentioned herein, it has been demonstrated that RNAi is a powerful tool to inhibit cellular gene expression. These studies enabled the development of peptides that facilitate binding of siRNA and simultaneously foster targeting specific receptors^10,11^ beyond the blood-brain barrier. A major deterrant that arose from these carriers is the presence of cationic charge that led to non-specific biodistribution of heterogeneously formed particles^12,13^. This aspect was addressed and overcome through studies using versatile approaches by the use of designed fusion proteins, where association with nucleic acids such as siRNA being conformation-dependent, did not compromise the function of targeting or alter the physiological stability when delivered *in vivo*. Towards this goal, observations from our lab established that engineered recombinant proteins have the potential to bring about specific and efficient gene silencing for cancer therapy that was experimentally proven in cancer xenograft mouse models^14^. The resultant association of these fusion proteins with siRNA led to physiological stability and functionality of the complex upon *in vivo* delivery. This is also backed by a recent proof-of-concept study^15^, where TARBP-BTP, an engineered fusion construct enabled efficient transport of siRNA across the BBB to the brain parenchyma resulting in consequential knockdown of BACE1 *in vivo*. The study demonstrated delivery of siRNA, mediated by engineered TARBP-BTP fusion protein, which led to specific knockdown of β-secretase (BACE1) significantly in cortical and sub cortical regions of the brain in both AβPP-PS1 and wild type C57BL/6 mice. In the current report, we extended the study further in AβPP-PS1 mice with the idea of devising a treatment procedure where long-term administration of siRNA, through BACE1 knockdown mediated by TARBP-BTP, would serve as a reliable strategy to decrease load of amyloid plaques in the mouse brain. Our data substantiates the approach and potential of TARBP-BTP as an efficient and promising platform to foster efficient RNAi by the delivery of BACE1 siRNA as an inhibitor of a key enzyme implicated in AD progression.

## Materials and Methods

Monoclonal Anti-Glyceraldehyde-3-Phosphate Dehydrogenase (GAPDH) antibody (MAB374) was purchased from Millipore. Anti Aβ antibody (ab2539), Anti BACE1 antibody (ab108394), Goat Anti-Rabbit IgG H&L (Alexa Fluor^®^ 647) (ab181474) were procured from Abcam. Congo Red (234610) was from Calbiochem. Ni Sepharose 6 Fast Flow was purchased from GE Healthcare. All other reagents used were of highest purity.

### Animals

Both AβPP-PS1 and wildtype control mice were maintained at an ambient temperature of 22°C in sterile individually ventilated containers, and 12/12 h dark/light cycles. The mice were given free access to water and food. All the experiments were conducted according to the guidelines of the Institute’s Animal Ethics Committee (IAEC) vide approved project # IAEC 76A/ 2016.

### TARBP-BTP purification

Recombinant protein TARBP-BTP was purified as described earlier^15^. Briefly, the protein expressed in *Escherichia coli* BL21(DE3) was purified using Ni-NTA affinity chromatography under denaturing conditions using urea. Cells were lysed in a buffer containing 6M urea, 50mM sodium phosphate buffer pH 7.4, 300 mM NaCl, 10 mM Tris, and 1mM PMSF followed by sonication. The lysate was clarified by centrifuging at high speed and the supernatant was added to Ni-NTA sepharose matrix. The matrix, loaded onto a gravity column, was washed with 0.1 % Triton X-114 in lysis buffer at 4°C to remove bacterial endotoxins^16^. Bound his-tag proteins were refolded by gradually reducing urea concentration and eluted using sodium phosphate buffer containing 300mM imidazole (pH 7.4) The eluted protein was dialysed against PBS (pH 7.4) overnight. TARBP-BTP:siRNA complex was prepared as described previously^15^. siRNA gradually diluted in PBS buffer was slowly added to an equal volume of TARBP-BTP taken at 5-fold molar excess.

### Experimental design

The study was conducted in AβPP-PS1 double transgenic mice. The complex was prepared and stored on ice 15 min prior to injection. Two groups of genotyped AβPP-PS1 of 8-9 month old mice received either TARBP-BTP or TARBP-BTP: BACE1 siRNA. A third group, which is the age matched non-transgenic mice, received normal saline. Mice were injected intravenously with the complex, prepared as described in the earlier section, via the tail vein every 4^th^ day. Totally seven injections were administered over equally spaced intervals for 28 days. Following treatment, mice were subjected to Morris Water Maze (MWM) Task as described in the next section for memory analysis, following which mice were euthanized using isoflurane, and quantified for Aβ plaques and histopathology analysis.

### Morris water maze task

Following treatment described in the preceeding section, mice were subjected to Morris water maze task to determine their behavioural ability of spatial learning and memory^17^. The task was carried out in a circular tank (diameter 150 cm) containing a platform, fixed in one quadrant, hidden 1 cm below the water level. The tank is virtually divided into 4 quadrants, and cards having specific patterns were kept as cues on the wall of the tank. The movement paths of mice were video recorded using a camera, and analysed using Ethovision software. During training period (1^st^ to 4^th^ day), mice were allowed to explore for 90 s, to find the platform by consecutively dropping them in the 3 quadrants. During this period the escape latency i.e. the time taken to reach the platform was monitored. The memory test was carried out on the 7^th^ and 8^th^ day following four days of training. For the first test, mice were dropped in the quadrant farthest from the platform, and the latency to reach the platform was recorded. For the memory test on second day, the platform was removed, and the time spent by the mice in that quadrant was recorded.

### Tissue preparation

At the end of the MWM task, blood was collected retro-orbitally from mice that were then euthanized by cervical dislocation. The different organs including brain were rapidly dissected and washed in PBS. The brain was sliced laterally into two halves. The first half was dissected to isolate various brain regions including cerebral cortex and hippocampus using a brain matrix and snap frozen in liquid nitrogen. The other half was used for immunohistochemistry analysis. For histochemistry examination, the brain tissue was fixed in 4% PFA for 20 h followed by transferring the tissues into 30% sucrose solution, and then frozen in OCT media and cryosectioned to 20μm slices. These free floating coronal sections collected in PBS, were mounted onto Fisher Probe-on Plus slides and air-dried overnight. To eliminate handling errors and to ensure consistant staining, sections from all three groups were mounted to the same slide. Parellel sections were subjected to both immunohistochemistry and Congo Red staining as described below.

### Aβ immunostaining

Aβ antibody recognizing epitope consisting of amino acids 1-14 of Aβ peptide was used in this study. Mounted cryosections were washed with PBS followed by permeabilization with 0.1% Tween 20. The sections were washed with PBS and blocked with 5% serum in PBST. This was followed by incubation with primary antibody (1:200 dilution in PBST) overnight at 4°C, then washed thrice and treated with secondary antibody at 4°C for 1 h. The slides were washed again, counterstained with Hoechst 33342, and mounted using vectashield mounting media. The images were acquired using Leica SP5 AOBS confocal microscope with a 10X dry objective and quantitated by NIH-ImageJ.

### Congo red staining for amyloid plaques

The coronal brain sections of mice corresponding to various treatment groups were subjected to Congo red staining to visualize amyloid plaques. The protocol has been described in detail elsewhere^18^. Briefly, the slide mounted, overnight air-dried brain cryosections were hydrated, followed by treatment with 80% ethanol saturated with NaCl for 20 min. The slides were subsequently treated with 0.2% Congo red solution prepared in 80% ethanol saturated NaCl solution. Prior to the treatment, both solutions were rendered alkaline by the addition of 10 mM NaOH. After 30 min exposure to Congo red solution, slides were washed in 95% and 100% ethanol and cleared in xylene. The coverslips were mounted using DPX mountant, and imaged using Axioplan upright microscope (Zeiss). Congo red-positive amyloid plaques were quantitated using NIH-ImageJ software. The images were segmented by HSB method to differentiate background and the Congo red positive pixels, and the area of these pixels were quantified.

### Histopathology

Tissues were cryosectioned to 12μm thickness and subjected to Hematoxylin and Eosin staining. Briefly, sections were air dried for 15 min and dipped in isopropanol solution for 10 min. The sections were then washed in distilled water and stained with Harris hematoxylin solution for 2 min in a dark place. Following washing with distilled water, slides were stained with eosin and immediately washed in distilled water. The slides were further dehydrated by treating with 50%, 70%, 95% and 100% ethanol. Finally, the slides were cleared in xylene and mounted using DPX mountant and visualized using Axioplan upright microscope (Zeiss).

### Immunoblotting

Hippocampal region of the brain of mice was lysed using a handheld homogeniser in Radio Immunoprecipitation Assay buffer (50 mM Tris–HCl pH 8.0, 150 mM NaCl, 2 mM EDTA, 1% NP-40, 0.5% sodium deoxycholate, 0.1% SDS) containing protease cocktail inhibitor. The lysate was clarified by centrifugation, and the supernatant was used for immunoblotting. The total protein content of the lysate was assessed by modified Lowry’s method^19^, and resolved on a SDS PAGE (12%) gel by loading 25μg of protein per lane. Protein was subsequently transferred on to nitrocellulose membrane and probed with antibodies for BACE1 and GAPDH. Blots were treated with the corresponding secondary antibody tagged with HRP and visualized using Pierce^®^ ECL Western Blotting Substrate (Thermo Scientific).

## Results and discussion

According to the amyloid cascade hypothesis, overproduction of Aβ peptide increases the formation of senile plaques, which in turn leads to synaptic dysfunction and neurodegeneration^1,2^. One promising approach to treat Alzheimer’s disease is to block the secretase enzymes which are responsible for the production of toxic Aβ peptide through cleavage of the amyloid precursor protein (APP). Inhibiting gamma secretase has a positive effect in reducing Aβ levels in the brain; however, it is associated with off target complications as gamma secretase has an important role in notch processing. β-Secretase is considered a safer target for AD as it is not critical in the normal functioning of the brain. However, new studies indicate that BACE1 may have other roles in axonal guidance, myelination etc^20^. Nevertheless, being a pharmaceutical target for AD, a variety of drugs from small molecules to designed antibodies targeting beta secretases have been tested^21–23^.

RNA interference, a post transcriptional regulatory mechanism found in higher organisms render the possibility of precisely modifying disease related genes, and where design of suitable carriers of siRNA leads to highly specific knockdown of the target gene. In an important work carried out by Singer et.al in 2005, lentiviral vectors expressing BACE1 siRNA upon stereotactic injection were shown to reduce the plaque levels and improve spatial learning and memory in APP mice^24^. However, the lack of suitable carriers for non invasive administration of siRNA hindered the translational potential of this strategy. Noting the efficacy of TARBP-BTP from our earlier study, in this report, we validated the delivery efficacy in a long-term concept study in AβPP-PS1 mouse models of AD. These mice were genotyped using gene specific primers (PS1) prior to treatment. The overall experimental strategy targeting BACE1 is depicted as shown in the schematic (**Fig. 1**).

**Fig.1.**
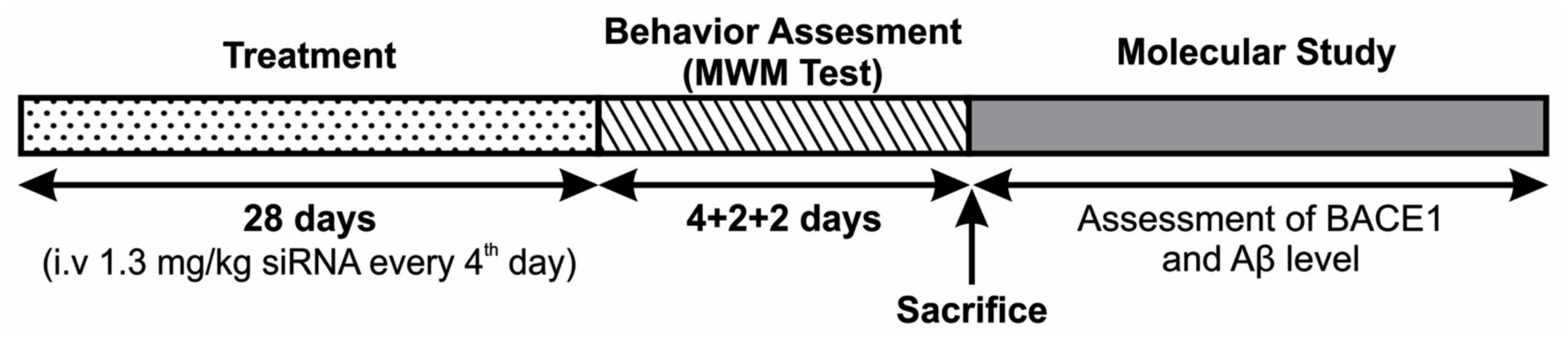
Experimental strategy and work flow.

The delivery complex, TARBP-BTP: BACE1 siRNA, was injected into the tail vein of mice as described in methods. Two groups of AβPP-PS1 mice (8-9 months) received either TARBP-BTP (vehicle) as a control group and TARBP-BTP:BACE1-siRNA complex (vehicle: siRNA) as the treated group. Mice were treated by the administration of TARBP-BTP or TARBP-BTP:BACE1-siRNA complex at an optimized dosage spanning a period of one month. Age matched non-transgenic mice that received normal saline constituted the third group. Based on our earlier gene knockdown studies and previously reported studies on the kinetics of siRNA mediated knockdown^25^, we planned a treatment strategy wherein the complex was injected every 4^th^ day, for a period of 28 days, to keep the BACE1 mRNA and protein levels consistently low.

Following treatment, we first assessed whether functional delivery of BACE1-siRNA has improved the behavioural function of spatial learning and memory by subjecting mice to Morris Water Maze test. As depicted (**Fig. 2a**), the escape latency of mice in all groups improved with training. However, there was a nonsignificant improvement in the escape latency of treated groups compared to the transgenic ones. Also, during Test 1 (**Fig. 2b**), none of the control transgenic mice could locate the platform in the given 90 s, while treated mice, on an average, took 48.8 s to find the platform (p=0.064). In Test 2, treated mice spent a significantly higher time i.e. 29.3 s in the target quadrant while the control group spent only 6.7 s (**Fig. 2c**). Both Test 1 and 2, revealed significant improvement in memory of treated mice, where mice showed reduced escape latency in Test 1. During Test 2, mice spent more time than the control group in the target quadrant. This suggests that the delivery of BACE1-siRNA mediated by TARBP-BTP has the potential to mediate target specific effects and improve cognitive functions in AD mice. Reduction in BACE1 levels leads to a considerable depletion in Aβ plaque levels, which otherwise will lead to cognitive impairments.

**Fig. 2.**
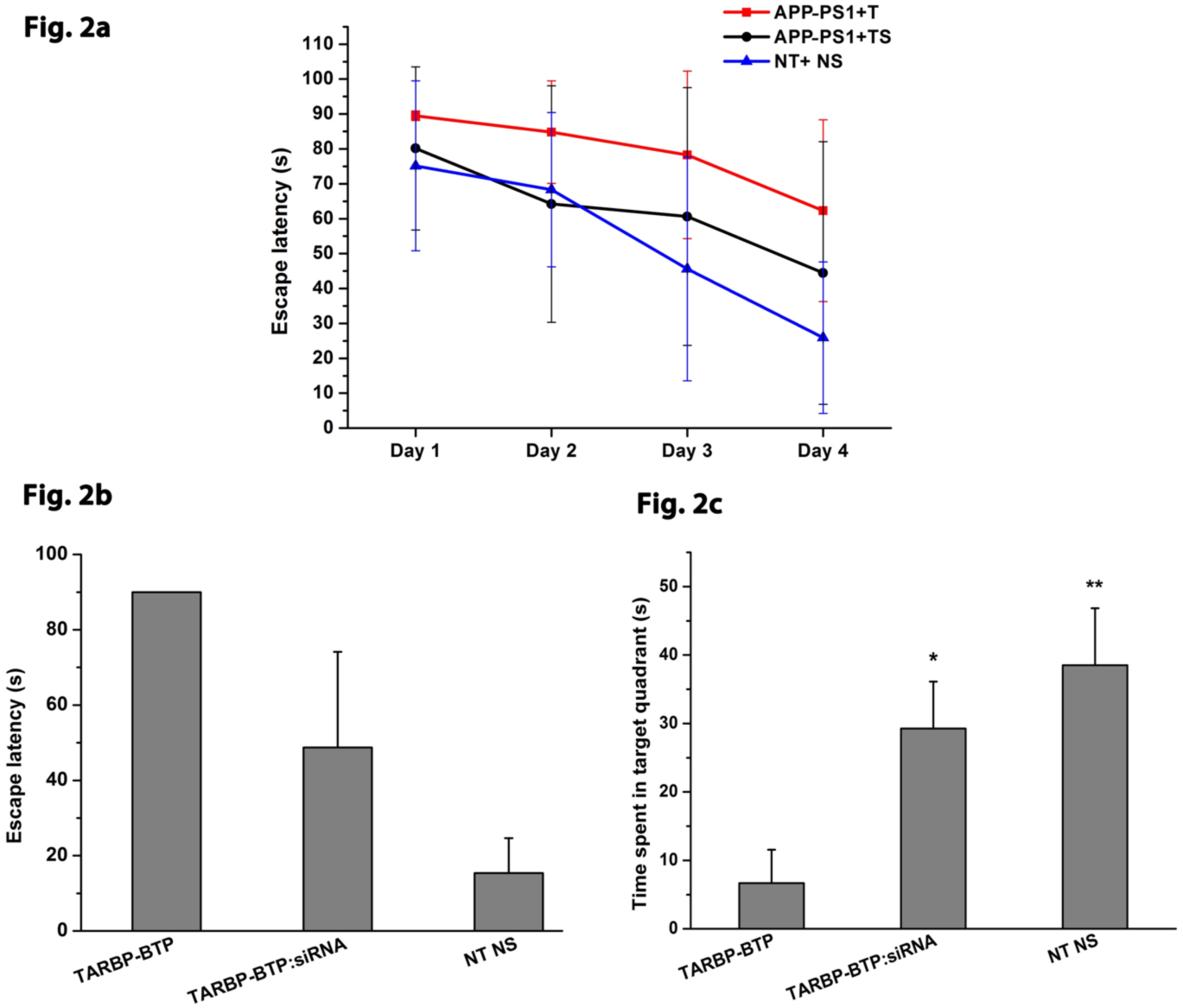
Morris Water Maze task (MWM) performed as described in methods, a) Learning curve depicting the time taken by mice to find the hidden platform during training. T= TARBP-BTP, TS= TARBP-BTP:BACE1-siRNA, NT+NS= Non-transgenic + normal saline. b) Escape latency of the mice group in Test 1, and c) Time spent by mice in the target quadrant during Test 2. Except NT-NS, other two groups were AβPP-PSl mice (n = 3) for TARBP-BTP, NT NS groups, and TARBP-BTP: BACE1-siRNA (n=4) group. Error bar indicates standard deviation, ** indicates p<0.01; * p<0.05 compared with vehicle only.

To assess whether the improvement in memory observed in the treated mice is a direct consequence of reduction in Aβ plaque, we carried out immunohistochemistry analysis of brain tissues using anti-Aβ antibody. As depicted, we observed a marked reduction in the plaque load in the treated group compared to the control (**Fig. 3a**). We also confirmed the observed reduction by staining brain sections with Congo Red, a dye that binds to the amyloid fibrils. As depicted (**Fig. 4a**), Congo red positive Aβ plaques were also significantly less in TARBP-BTP:BACE1-siRNA-treated mice. Quantitative analysis of Aβ levels by both techniques conclusively showed significantly less plaque levels in the treated group compared to controls (**Fig. 3b** and **4b**). Also, analysis of BACE1 protein expression levels in these mice showed considerable reduction in the treated groups compared to vehicle treated controls (**Fig. 4c**). In contrast, higher expression of BACE1 in control AβPP-PS1 compared to non transgenic mice is expected. Our results clearly indicate that treatment of mice with BACE1-siRNA, for a period of one month, led to consistent reduction in BACE1 levels leading to reduced cleavage of APP molecule. This leads to likely reduction in Aβ peptide levels which in turn results in the improvement in the memory.

**Fig. 3.**
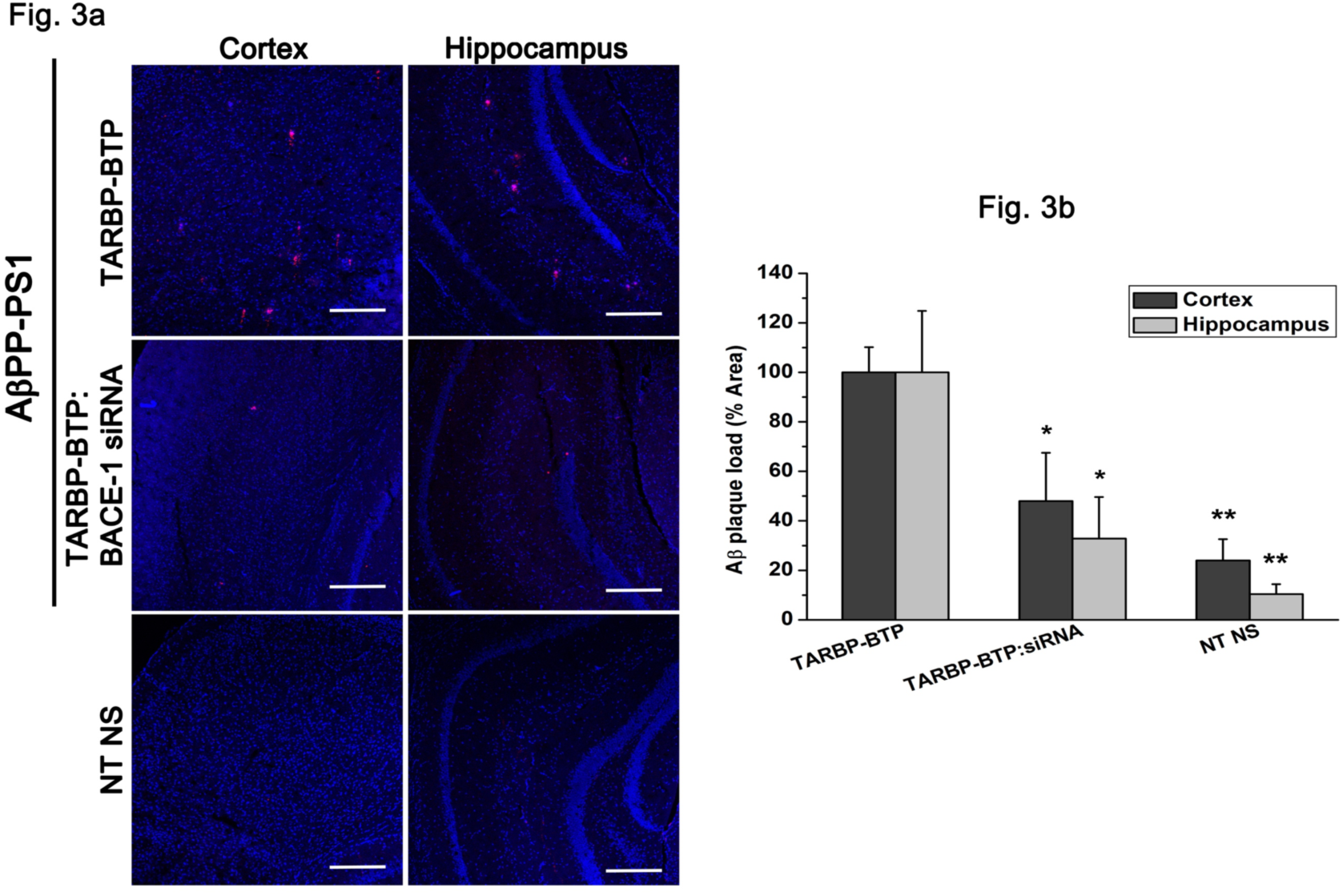
a) Immunohistochemical analysis of Aβ plaque load in coronal sections of mice hippocampus and cerebral cortex visualized by Aβ antibody staining (red) and counterstained with Hoechst 33342 (blue). Images were obtained using Leica SP5 confocal microscope, Mag 10X, scale bar =200μm. b) Quantitation of Aβ plaque area by imageJ. Values were normalized to the respective controls (n=2) for TARBP-BTP, and NT-NS (n = 2) for TARBP-BTP:BACE1-siRNA. Error bar indicates standard deviation, ** indicates p<0.01; * p<0.05 compared with vehicle only.

**Fig. 4.**
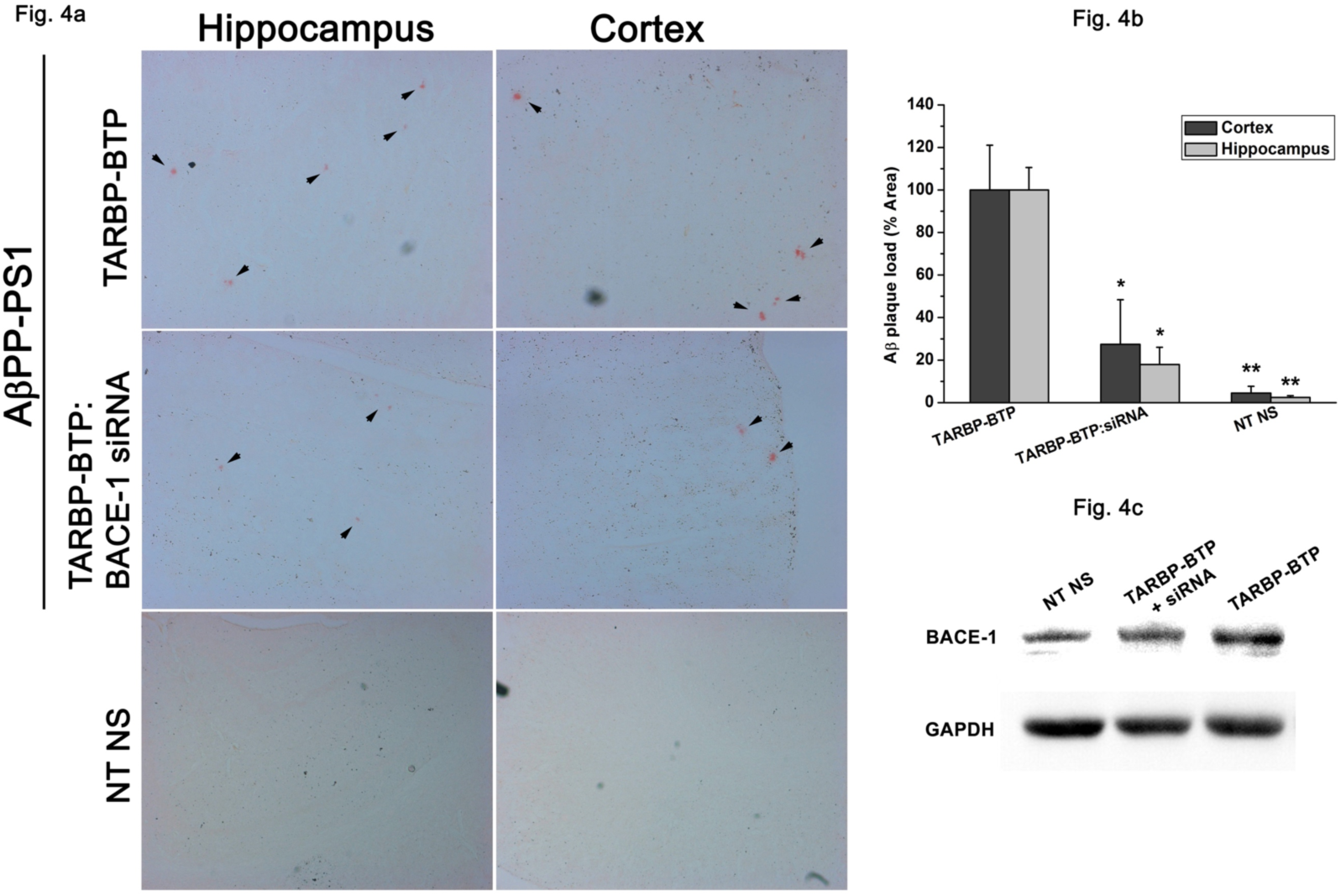
a) Visulisation of of Aβ plaques using Congo Red staining approach in coronal sections of hippocampas and cerebral cortex in mice. Arrow heads indicate Congo red-positive Aβ plaques. Magnificaion 10X. b) ImageJ quantitation of % area of Congo red pixels in both cerebral cortex and hippocampus. The values were normalized to respective controls (n=2) for all groups. c) Western blot of representative hippocampal tissue lysates from the three groups probed for BACE1 expression, GAPDH served as an internal control. Error bar indicates standard deviation, ** indicates p<0.01; * p<0.05 compared with vehicle only.

Safety and nontoxic nature are prime attributes of peptide-based biomaterials which holds potential for usage of protocols as described here eventually towards human therapy. Interestingly, repeated dose of TARBP-BTP:BACE1-siRNA complex did not result in any visible toxicity, adverse effects or behavioural changes in mice. We also investigated the possibility of inflammation and other toxicity by examining the vital organs such as spleen, liver, kidney, heart and lungs of these mice through histopathological analysis. Hematoxylin and Eosin staining confirmed that there was no inflammation, periportal infilitration of leukocytes or tissue necrosis (**Fig. 5**). These results indicated the likely non-toxic nature of the delivered complex. However, possible longterm effects of the complex was not investigated by other methods during the course of this investigation.

**Fig. 5.**
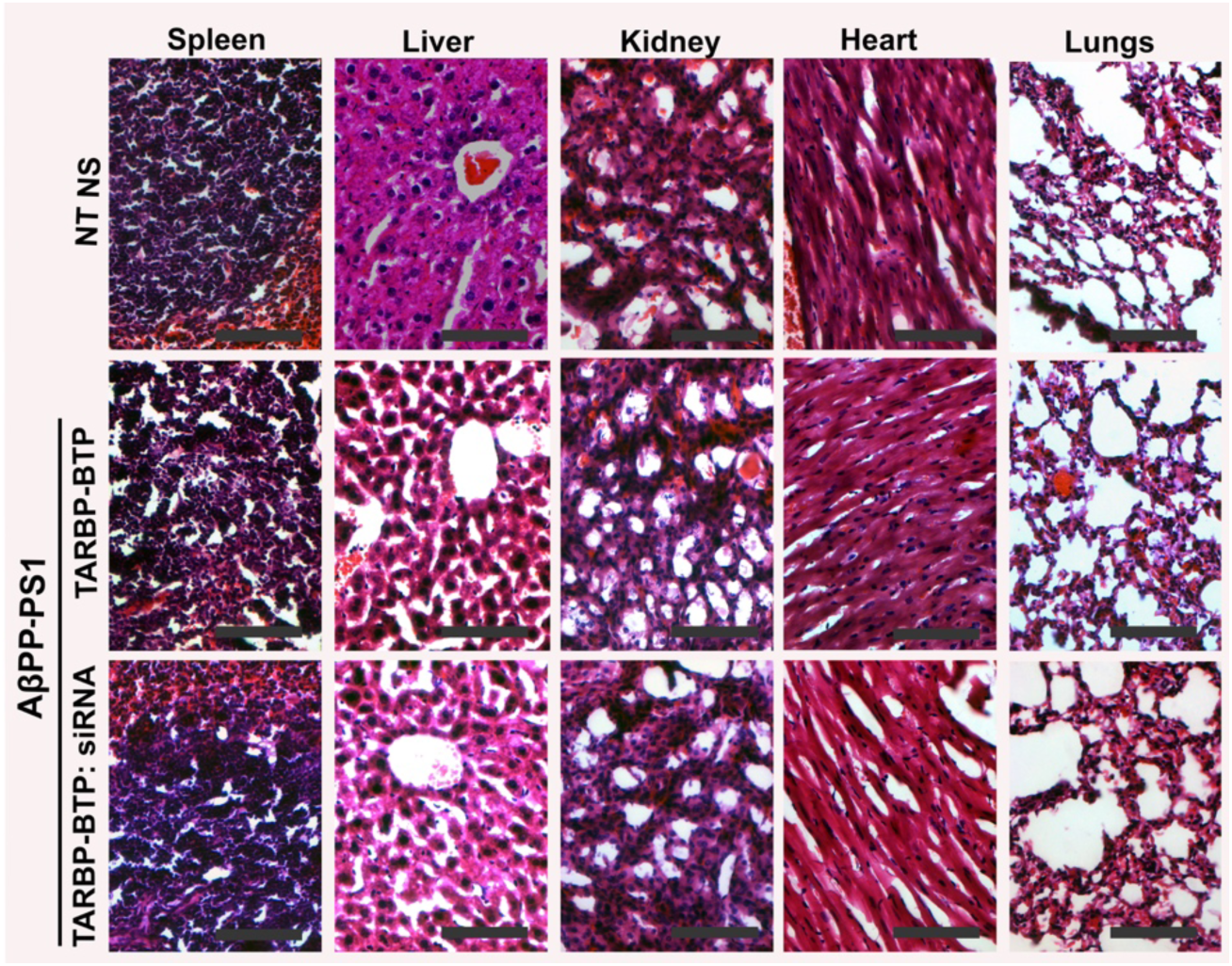
Representative Hematoxylin and Eosin stainied micrographs of the major organs. Magnification 20X, Scale bar=100 μM.

Our preceding report indicated the advantageous benefits of TARBP-BTP fusion protein that mediates target-specific delivery of therapeutic siRNA to brain tissues in AβPP-PS1 mouse model of AD^15^, leading to a promising therapeutic approach towards anti-amyloid therapy. The present study clearly validates the potential of TARBP-BTP fusion protein as a delivery platform in delivering RNAi-based interventions for AD. It is evident that functional knockdown of BACE1 leads to significant reduction in Aβ resulting in reversal of cognitive defects. In future, other parameters like dose escalation, dose response, usage of chemically modified siRNA for higher stability need to be explored to unravel the full potential, mandating extensive study in this direction. The study furthermore illustrates the suitability of the carrier targeting other neurological disorders in addition to exploring largely unmet therapeutic needs and facilitating basic neuroscience research to realise the potential application of the strategy in gaining a deeper insight into the mechanism of diseases affecting the CNS. Importantly, the carrier enables synergistic delivery of a mixture of siRNAs to target key genes implicated in early onset of neurodegenerative disorders, an area of research that remains to be explored.

## Conclusions

The concept study demonstrates TARBP-BTP fusion protein as a promising carrier of siRNA in potentially therapeutic quantities for AD intervention. The strategy described here will, in addition, have the capacity to advance basic and translational science through studies targeting neurodegenerative disorders.

## Acknowledgements

The work was supported by BRNS (DAE) Grant No. 37(1)/14/51/2014-BRNS. The authors thank Sairam for diligent help with animal dissections, and Nandini Rangaraj for help with confocal microscopy. We acknowledge Durga Jeyalakshmi Srinivasan’s inputs to the progress of this work, and timely assistance from Haritha Nunna with the *in vivo* experiments.

## References

1. Perl, D. P. Neuropathology of Alzheimer’s disease. Mount Sinai Journal of Medicine 77, 32–42 (2010).

2. Citron, M. Alzheimer’s disease: strategies for disease modification. Nat. Rev. Drug Discov. 9, 387–398 (2010).

3. Selkoe, D. J. & Schenk, D. Alzheimer’s disease: molecular understanding predicts amyloid-based therapeutics. Annu. Rev. Pharmacol. Toxicol. 43, 545–84 (2003).

4. Walsh, D. M. & Selkoe, D. J. Abeta oligomers - A decade of discovery. J. Neurochem. 101, 1172–1184 (2007).

5. Neuwelt, E. et al. Strategies to advance translational research into brain barriers. Lancet Neurol. 7, 84–96 (2008).

6. Hawkins, B. T. & Davis, T. P. The Blood-Brain Barrier / Neurovascular Unit in Health and Disease. Pharmacol. Rev. 57, 173–185 (2005).

7. Pardridge, W. M. The blood-brain barrier: bottleneck in brain drug development. NeuroRx 2, 3–14 (2005).

8. Sankaranarayanan, S. et al. In Vivo β-Secretase 1 Inhibition Leads to Brain Aβ Lowering and Increased β-Secretase Processing of Amyloid Precursor Protein without Effect on Neuregulin-1. Pharmacology 324, 957–969 (2008).

9. Citron, M. β-Secretase inhibition for the treatment of Alzheimer’s disease - Promise and challenge. Trends Pharmacol. Sci. 25, 92–97 (2004).

10. El-Andaloussi, S., Johansson, H. J., Holm, T. & Langel, Ü. A Novel Cell-penetrating Peptide, M918, for Efficient Delivery of Proteins and Peptide Nucleic Acids. Mol. Ther. 15, 1820–1826 (2007).

11. Kumar, P. et al. Transvascular delivery of small interfering RNA to the central nervous system. Nature 448, 39–43 (2007).

12. Schneider, B. et al. Targeted siRNA Delivery and mRNA Knockdown Mediated by Bispecific Digoxigenin-binding Antibodies. Mol. Ther. Nucleic Acids 1, e46 (2012).

13. Fiedel, B. A, Rent, R., Myhrman, R. & Gewurz, H. Complement activation by interaction of polyanions and polycations. Immunology 30, 161–9 (1976).

14. Dar, G. H., Gopal, V. & Rao, M. Conformation-dependent binding and tumor-targeted delivery of siRNA by a designed TRBP2: Affibody fusion protein. Nanomedicine Nanotechnology, Biol. Med. 11, 1455–1466 (2015).

15. Haroon, M. M. et al. A designed recombinant fusion protein for targeted delivery of siRNA to the mouse brain. J. Control. Release 228, 120–131 (2016).

16. Reichelt, P., Schwarz, C. & Donzeau, M. Single step protocol to purify recombinant proteins with low endotoxin contents. Protein Expr. Purif. 46, 483–488 (2006).

17. Vorhees, C. V & Williams, M. T. Morris water maze: procedures for assessing spatial and related forms of learning and memory. Nat Protoc 1, 848–858 (2006).

18. Wilcock, D. M., Gordon, M. N. & Morgan, D. Quantification of cerebral amyloid angiopathy and parenchymal amyloid plaques with Congo red histochemical stain. Nat. Protoc. 1, 1591–1595 (2006).

19. Markwell, M. A. K., Haas, S. M., Bieber, L. L. & Tolbert, Ne. A modification of the Lowry procedure to simplify protein determination in membrane and lipoprotein samples. Anal. Biochem. 87, 206–210 (1978).

20. Barão, S., Moechars, D., Lichtenthaler, S. F. & De Strooper, B. BACE1 Physiological Functions May Limit Its Use as Therapeutic Target for Alzheimer’s Disease. Trends in Neurosciences 39, 158–169 (2016).

21. Vassar, R. BACE1 inhibitor drugs in clinical trials for Alzheimer’s disease. Alzheimers. Res. Ther. 6, 89 (2014).

22. Atwal, J. K. et al. A therapeutic antibody targeting BACE1 inhibits amyloid-β production in vivo. Sci. Transl. Med. 3, 84ra43 (2011).

23. Kennedy, M. E. et al. The BACE1 inhibitor verubecestat (MK-8931) reduces CNS β-amyloid in animal models and in Alzheimer’s disease patients. 1–14 (2016). doi:10.1126/scitranslmed.aad9704

24. Singer, O. et al. Targeting BACE1 with siRNAs ameliorates Alzheimer disease neuropathology in a transgenic model. Nat. Neurosci. 8, 1343–1349 (2005).

25. Bartlett, D. W. & Davis, M. E. Insights into the kinetics of siRNA-mediated gene silencing from live-cell and live-animal bioluminescent imaging. Nucleic Acids Res. 34, 322–333 (2006).

